# High genetic diversity of *Plasmodium falciparum* in the low transmission setting of the Kingdom of Eswatini

**DOI:** 10.1101/522896

**Authors:** Michelle E. Roh, Sofonias K. Tessema, Maxwell Murphy, Nomcebo Nhlabathi, Nomcebo Mkhonta, Sibonakaliso Vilakati, Nyasatu Ntshalintshali, Manik Saini, Gugu Maphalala, Anna Chen, Jordan Wilheim, Lisa Prach, Roly Gosling, Simon Kunene, Michelle Hsiang, Bryan Greenhouse

## Abstract

**ABSTRACT:** *Background:* To better understand transmission dynamics, we characterized *Plasmodium falciparum* (*Pf*) genetic diversity in Eswatini, where transmission is low and sustained by importation.

*Methods:* 26 *Pf* microsatellites were genotyped in 66% of all confirmed cases from 2014-2016 (n=582). Population and within-host diversity were used to characterize differences between imported and locally-acquired infections, as determined by travel history. Logistic regression was used to assess the added value of diversity metrics to classify imported and local infections beyond epidemiology data alone.

*Results:* The parasite population in Eswatini was highly diverse (H_E_=0.75) and complex, with 67% polyclonal infections, a mean MOI of 2.2, and mean F_WS_ of 0.84. Imported cases had comparable diversity to local cases, but exhibited higher MOI (2.4 versus 2.0; p=0.004) and lower mean F_WS_ (0.82 vs. 0.85; p=0.03). Addition of MOI and F_WS_ to multivariate analyses did not increase discrimination between imported and local infections.

*Discussion:* In contrast to the commonly held perception that *Pf* diversity declines with decreasing transmission intensity, isolates from Eswatini exhibited high parasite diversity consistent with high rates of malaria importation and limited local transmission. Estimates of malaria transmission intensity from genetic data need to consider the effect of importation, especially as countries near elimination.

## INTRODUCTION

The Kingdom of Eswatini, formerly known as the Kingdom of Swaziland, has made significant progress in reducing its malaria burden and is on track to be one of the first sub-Saharan African countries to eliminate malaria. From 2000-2010, Eswatini reduced its malaria cases by 94% [1] and has since sustained very low malaria transmission, reporting an average of 1.5 cases per 1000 over the period of 2010-2016 [2]. A key challenge in eliminating malaria in Eswatini has been cross-border migration from neighboring, high transmission countries, which has resulted in malaria importation and sporadic onward transmission in malaria receptive areas [3-5]. Between 2014 and 2016, a total of 55% of infections were classified as imported [6], mainly from Mozambique (90%) [4, 7].

To complement traditional approaches to surveillance, genotyping of *P. falciparum* has become a popular approach to understanding the underlying malaria transmission dynamics [8-10]. As transmission declines, recombination between genetically distinct clones likely reduces, leading to decreases in genetic diversity, effective population size and fragmented populations [11]. This relationship has been proposed to be potentially useful as a “genomic thermometer” to gauge changes in malaria transmission resulting from intensified malaria control and elimination efforts [8, 12, 13]. This “genomic thermometer” framework has been based on numerous studies that have shown that *P. falciparum* parasite populations generally have higher population and within-host diversity (i.e. higher multiplicity of infection or MOI) in high transmission settings of sub-Saharan Africa than in lower transmission settings such as Southeast Asia and Latin America [8, 11, 14-22]. However, studies of *P. falciparum* parasite diversity in low transmission settings of sub-Saharan Africa are limited, and in these settings, importation of malaria from neighboring, high transmission countries may complicate the relationship between transmission intensity and genetic diversity.

Here, we evaluate the *P. falciparum* parasite population in Eswatini, a low transmission country challenged by malaria importation from bordering countries with high malaria transmission. This study evaluates how well parasite diversity aligns with this “genomic thermometer” framework in Eswatini and evaluates the utility of parasite diversity metrics to distinguish between imported and locally-acquired infections.

## METHODS

### Study site

Eswatini is a small, landlocked country in Southern Africa bordered by South Africa and Mozambique, with an estimated population size of 1.3 million [23]. Approximately 30% of the population live in the eastern half of the country, which is still receptive to malaria. In these areas, malaria transmission is unstable and highly dependent on rainfall and cross-border malaria importation [5, 7, 24].

### National malaria field surveillance and initial molecular testing

Malaria is a notifiable disease in Eswatini and all confirmed cases are reported to the national malaria surveillance system [25]. Cases are confirmed by either microscopy or rapid diagnostic test (RDT) and followed up for investigation, whereby demographics, household GPS coordinates, and travel history from the prior eight weeks were collected. Based on investigation data, cases are classified as imported if they reported travel to a higher endemic country in the past eight weeks. In this study, investigators imposed a further restriction by reclassifying imported cases as local if travel was in the week prior to case detection to account for the incubation period of *P. falciparum*. If the case resided in a malaria-receptive area, reactive case detection (RACD), or focal “test and treat” using RDTs, is conducted around the household of the “index case” (i.e. the case that triggered the RACD event [26]). Similar epidemiological data are collected from individuals during RACD events and dried blood spots (DBS) are collected from both index cases and individuals screened during RACD.

DBS samples underwent molecular testing by loop-mediated isothermal amplification (LAMP) using previously described methods [27]. A second DBS from all index cases and LAMP-positive individuals from RACD was collected for subsequent quantitative polyermase chain reaction (PCR) testing at the University of California, San Francisco (UCSF) using the *varATS* method [28]. Samples were transported at room temperature and stored at −20°C. In both laboratories, DNA was extracted from DBS using the saponin Chelex method [29].

### Study population

Study eligibility included cases identified through Eswatini’s national malaria surveillance system between July 2014 and June 2016 (e.g. index cases and secondary cases identified through RACD). Cases had to have a DBS available for genotyping and a parasite density of ≥10 parasites/μL. Ethics approval was granted by the Eswatini Ethics Committee and the Committee on Human Research at the University of California, San Francisco.

### Microsatellite genotyping of P. falciparum

Samples were genotyped using microsatellite markers at 26 loci distributed across the *P. falciparum* genome [11, 30, 31] (Liu et al, in preparation). To minimize genotyping errors, strict thresholds were used to call alleles using a semi-supervised naïve Bayes classifier implemented in microSPAT software [32]. Capillary electrophoresis peaks were excluded if the probability of being a true allele was less than 95%. Samples that were genotyped at ≤13 loci were excluded.

### Characterization of population-level diversity

Population-level genetic diversity was assessed by expected heterozygosity (H_E_), the number of alleles per locus (i.e allelic richness), and the effective population size (N_e_). H_E_ is defined as the probability of randomly drawing a pair of different alleles from the allele pool. Values for H_E_ range from 0-1 (0 indicating no diversity and 1 indicating 100% of alleles are different). H_E_ was calculated for each locus using the formula 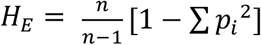, where *n*=the number of samples analyzed, *p_i_*=the allele frequency of the *i^th^* allele in the population. Mean H_E_ was calculated by taking the average of H_E_ across all loci. The number of unique alleles per locus, defined as A, is an alternative measure of population-level genetic diversity. The effective population size (N_e_), was estimated using the infinite alleles model (IAM) and the step-wise mutation model (SMM) as previously described [11]. Genetic differentiation between groups was assessed by comparing differences in H_E_ using the Hedrick’s G’_ST_ method [33].

To compare Eswatini’s *P. falciparum* genetic diversity relative to other malaria-endemic countries, a literature search of studies that used a similar subset of microsatellite markers was conducted and only studies that published expected heterozygosities at each loci were included. Estimates of population-level diversity from Eswatini were then recalculated using the same subset of microsatellite markers used in each comparative study and multiple alleles were scored if minor peaks were >25% of the height of the predominant allele.

### Characterization of within-host diversity

Within-host diversity was assessed by multiplicity of infection (MOI), the proportion of polyclonal infections and the within-host infection fixation index (F_WS_). MOI was defined as the second highest number of alleles observed in any one of the 26 loci that were genotyped. An infection was considered polyclonal if MOI was greater than one. The F_WS_ metric is a measure of within-host diversity that describes the relationship between the genetic diversity of an individual infection relative to the genetic diversity of the parasite population [34]. A low F_WS_ indicates high within-host diversity relative to the population (e.g. low risk of inbreeding). F_WS_ was calculated for each sample using the equation 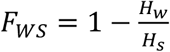, where *H_w_* = allele frequency of each unique allele found at a particular locus for each individual and *H_s_* = heterozygosity of the local parasite population. Within host allele proportions were estimated by relative capillary electrophoresis peak size. Amplification bias of different sized amplicons was corrected by estimating preferential bias as a function of amplicon size difference using known controls at known relative densities [32]. Using previously described F_WS_ thresholds [17, 35], samples with a F_WS_>0.95 were considered clonal infections and samples with a F_WS_ ≤0.70 were considered highly diverse infections.

### Predictive ability of MOI and FWS to discriminate between imported and local cases

Multivariate logistic regression was used to assess whether the addition of MOI and mean F_WS_ to epidemiologically parameterized models would predict whether a case was imported or locally-acquired. The base epidemiological model included predictors other than travel history that would be commonly used distinguish imported versus local cases, including: district of residence, season, whether a case was passively or reactively detected, age, gender, and occupation. Model improvement was tested using the log-likelihood chi-squared statistic and discriminative ability was assessed by the change in the area under the receiver operating characteristic (ROC) curve.

### Sensitivity analyses

Sensitivity analyses were conducted to assess whether results obtained from genotyped cases were representative of results that would have been obtained from all cases detected during the 2014-2016 period. Due to missing DBS samples or genotype failures, a proportion of cases were not genotyped (n=298) which may have resulted in a non-random sampling of the original case population (n=880). Comparisons between genotyped and non-genotyped cases revealed that these two groups significantly differed by several variables, including: case detection method (passive versus reactive surveillance), season, district, case classification (imported versus local), gender, and occupation (Supplementary Table 1). To account for this potential source of selection bias, we calculated a propensity score using a logistic regression model [36] which included the aforementioned demographic variables as predictors to estimate the probability of each case being genotyped. The inverse of these probabilities were used to re-weight within-host diversity measures obtained from genotyped individuals to represent the total case population. To create a propensity score model that would best predict the probability of being genotyped, statistical interactions were tested with each possible predictor combination and all interactions that had a p-value <0.10 were included in the model. Other than three non-genotyped cases that had a propensity score <0.17, density plots showed good overlap of the propensity scores between genotyped and non-genotyped groups (Supplementary Figure 1), suggesting reweighting by propensity scores was an appropriate approach to obtain generalizable estimates. None of the results were materially affected by this reweighting, therefore the primary results presented below all use the original, unweighted data.

### Statistical tests

All analyses were performed using Stata 14.0 (StataCorp, College Station, TX) and R (version 3.5.0; R Project for Statistical Computing; http://www.r-project.org/). Statistical comparisons were conducted using Pearson’s chi-squared test for categorical variables; and Student’s t-tests or Mann-Whitney tests for continuous variables, depending on the degree of normality of underlying distributions. The Spearman’s rank correlation (ρ) was used to assess the correlation between mean F_WS_ values and MOI. P-values less than 0.05 were considered statistically significant.

## RESULTS

### Characteristics of samples and genotyping

Between July 2014 and June 2016, 880 P. *falciparum* cases were identified through Eswatini’s national malaria surveillance program (Figures 1 and 2). Between 2014-2015 and 2015-2016, Eswatini experienced a 2.6-fold reduction in cases (from 635 to 245 cases). The majority of cases were detected through passive surveillance (82%) and the rest were identified through reactive case detection. Of the cases identified, 666 *P. falciparum* positive samples were available for genotyping. Twenty-six microsatellite markers were used for genotyping, and the mean success rate across all 26 markers was 86% (range=74-94). Of the 666 genotyped samples, 582 (87%) samples were genotyped successfully at >13 loci and included in the study. Sixty-six percent of genotyped samples had full coverage for all 26 loci.

**Figure 1.**
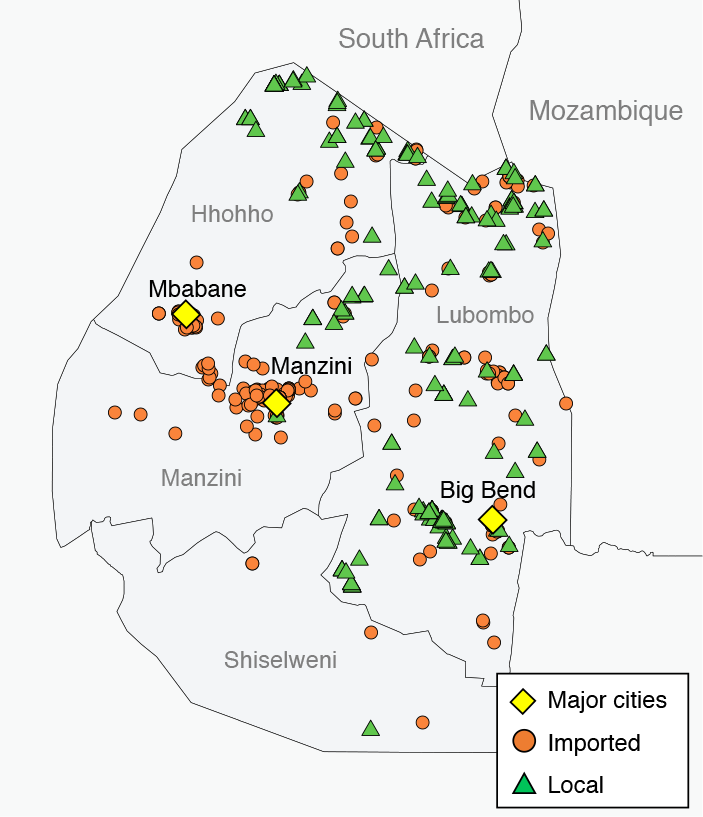
Map of Eswatini cases detected between July 2014 and June 2016. Orange circles represent imported cases and green triangles represent local cases. Yellow diamonds designate major Eswatini cities.

**Figure 2.**
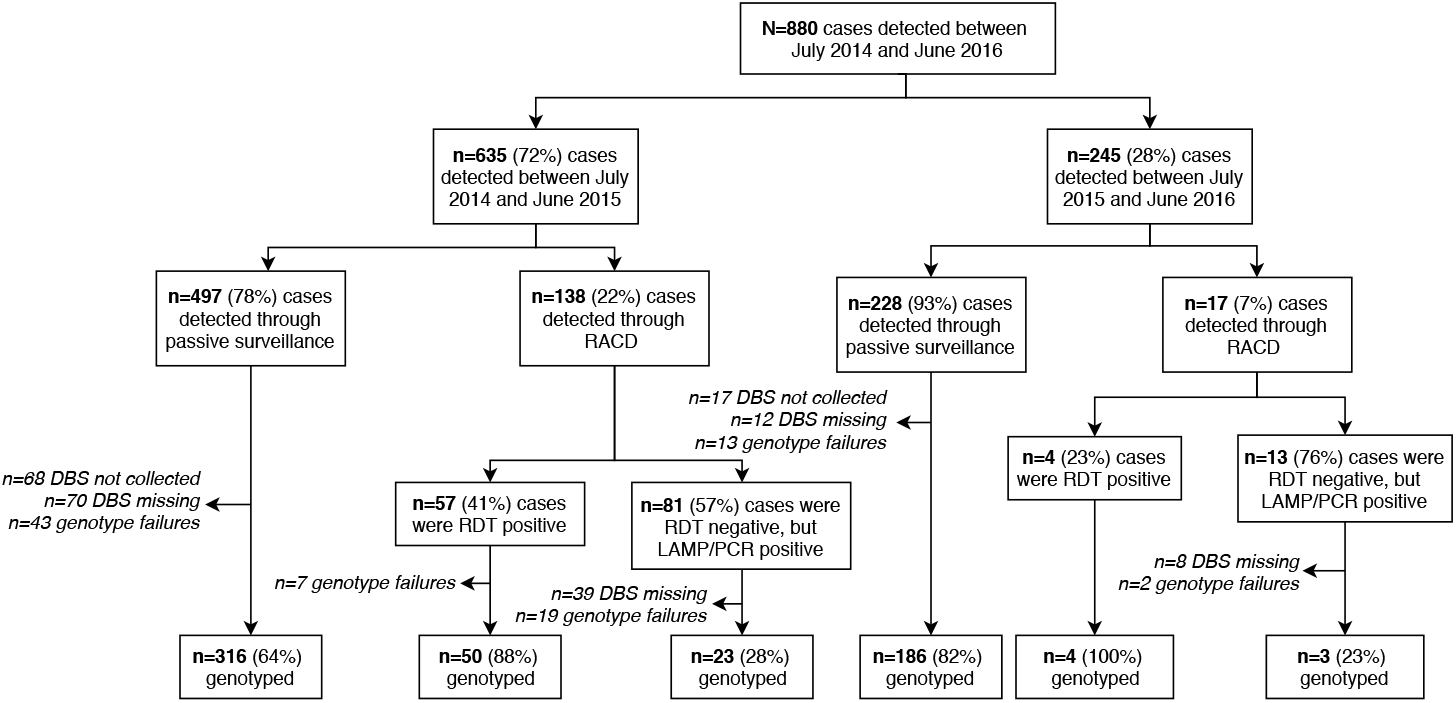
Flow chart of Eswatini samples collected and genotyped between July 2014 and June 2016. Abbreviations are as follows: DBS=dried blood spot; LAMP=loop-mediated isothermal amplification; PCR=polymerase chain reaction; RACD=reactive case detection; and RDT=rapid diagnostic test.

The characteristics of the sample population are presented in Supplementary Table 1. Sixty-two percent of genotyped cases (n=359) were classified as imported by National Malaria Program surveillance officers using self-reported travel history. Of the imported cases that had available country-level travel data (n=188), 93% reported traveling to Mozambique in the previous eight weeks, with a few cases reporting travel to either South Africa (n=5), Zambia (n=2), Zimbabwe (n=2), or Burundi (n=2). Two-hundred and fifteen infections were classified as local infections and only 12 of these were classified as intraported (i.e. imported from different areas within Eswatini [5]).

### Parasite populations in Eswatini are complex and diverse

Polyclonal infections were found in 66% of cases, with a mean multiplicity of infection (MOI) of 2.2 (range: 1-5) (Table 1). Within-host diversity was also assessed using the within-host infection fixation index (F_WS_) [34], whereby a lower F_WS_ demonstrates a higher within-host diversity [17]. Mean F_WS_ in the sample population was 0.84 (range: 0.44-1.00) and was negatively correlated with MOI (Supplementary Figure 2). Thirty-eight percent of cases had a F_WS_ value >0.95, a cut-off point used to indicate clonal infections, and 23% had a F_WS_≤0.70, a threshold for infections exhibiting high within-host diversity [17, 35]. At the population-level, parasite isolates from Eswatini were highly diverse (mean H_E_: 0.75; range: 0.52-0.93) (Table 2; Supplementary Table 2) and displayed high allelic richness (Mean A=15.4, SD: 7.1; range: 7-32) (Supplementary Table 3). The estimated effective population size (Ne) was 4717 [95% CI: 2027-10745] and 11792 [95% CI: 5067-26863] under IAM and SMM models, respectively.

Despite a 2.6-fold reduction in cases, within-host and population-level diversity were comparable between July to June seasons of 2014-2015 and 2015-2015 and displayed little genetic differentiation (G’_ST_=0.026). Similarly, no significant differences were observed between districts (G’_ST_=0.0068), except in Hhohho and Shiselweni districts, where the proportion of monoclonal infections were higher (50/115 or 44% and 3/5 or 60%, respectively) than in Manzini (71/235; 30%) and Lubombo districts (48/158; 30%) (p=0.032). No significant differences in within-host and population-level diversity and genetic differentiation (G’_ST_=0.003) were observed between passively versus reactively detected cases.

**Table 1.** Metrics of within-host diversity in Eswatini by group. Percentages are reported as % (95% confidence interval) and means are reported as mean ± standard deviation [range].

| Groups | % polyclonal | p-value<sup>b</sup> | Mean MOI | p-value<sup>b</sup> | Mean F<sub>WS</sub> | p-value<sup>b</sup> |
| --- | --- | --- | --- | --- | --- | --- |
| **Total (n=582)** | 67 (63-70) | -- | 2.2 ± 1.2 [1.0-7.0] | -- | 0.84 ± 0.15 [0.44-1.00] | -- |
| **Season** |  | 0.34 |  | 0.15 |  | 0.24 |
| June 2014-July 2015 (n=389) | 65 (60-70) | | 2.2 ± 1.2 [1.0-7.0] | | 0.84 ± 0.16 [0.44-1.00] | |
| June 2015-July 2016 (n=193) | 69 (63-76) | | 2.4 ± 1.2 [1.0-6.0] | | 0.83 ± 0.15 [0.49-1.00] | |
| **Districts** |  | 0.032 |  | 0.30 |  | 0.11 |
| Hhohho (n=115) | 57 (47-65) | | 2.1 ± 1.3 [1.0-7.0] | | 0.87 ± 0.15 [0.49-1.00] | |
| Lubombo (n=235) | 70 (64-75) | | 2.1 ± 1.0 [1.0-6.0] | | 0.83 ± 0.15 [0.47-1.00] | |
| Manzini (n=158) | 70 (62-76) | | 2.4 ± 1.3 [1.0-6.0] | | 0.83 ± 0.16 [0.44-1.00] | |
| Shiselweni (n=5) | 40 (12-77) | | 1.8 ± 1.3 [1.0-4.0] | | 0.86 ± 0.19 [0.58-1.00] | |
| **Passive versus reactive case detection** |  | 1.00 |  | 0.87 |  | 0.73 |
| Passive (n=502) | 67 (62-71) | | 2.2 ± 1.2 [1.0-7.0] | | 0.84 ± 0.15 | |
| Reactive (n=80) | 66 (55-76) | | 2.2 ± 1.2 [1.0-6.0] | | 0.83 ± 0.16 | |
| **Case classification** |  | 0.004 |  | 0.004 |  | 0.03 |
| Imported (n=359) | 71 (66-76) | | 2.4 ± 1.3 [1.0-7.0] | | 0.82 ± 0.16 [0.44-1.00] | |
| Local<sup>a</sup> (n=215) | 59 (52-65) | | 2.0 ± 1.1 [1.0-6.0] | | 0.85 ± 1.15 [0.47-1.00] | |

Note: MOI = multiplicity of infection; F<sub>WS</sub> = metric of within-host diversity relative to that of the population.
<sup>a</sup> Local infections include indigenous cases (n=203) and intraported (e.g. imported from different areas of the country) cases (n=12).
<sup>b</sup> All p-values were two-tailed and computed using the chi-squared test for comparing the proportion of polyclonal infections, Poisson regression for MOI, and simple linear regression for F<sub>WS</sub>.

**Table 2.** Metrics of population-level genetic diversity in Eswatini by group. All values are reported as mean ± standard deviation [range].

| Groups | Mean $H_E$ | Mean $A$ |
| --- | --- | --- |
| **Total (n=582)** | 0.75 ± 0.14 [0.52-0.93] | 15.4 ± 7.1 [7.0-32.0] |
| **Season** |  |  |
| June 2014-July 2015 (n=389) | 0.75 ± 0.14 [0.54-0.93] | 15.0 ± 6.9 [6.0-30.0] |
| June 2015-July 2016 (n=193) | 0.74 ± 0.16 [0.46-0.94] | 12.4 ± 6.6 [4.0-27.0] |
| **Districts** |  |  |
| Hhohho (n=115) | 0.75 ± 0.14 [0.50-0.92] | 12.0 ± 5.7 [4.0-22.0] |
| Lubombo (n=235) | 0.75 ± 0.14 [0.52-0.93] | 13.9 ± 6.5 [6.0-29.0] |
| Manzini (n=158) | 0.75 ± 0.15 [0.50-0.93] | 12.7 ± 6.3 [4.0-26.0] |
| Shiselweni (n=5) | 0.72 ± 0.19 [0.34-0.97] | 3.2 ± 1.3 [2.0-6.0] |
| **Passive versus reactive case detection** |  |  |
| Passive (n=502) | 0.75 ± 0.14 [0.52-0.93] | 15.3 ± 7.1 [7.0-31.0] |
| RACD (n=80) | 0.75 ± 0.14 [0.51-0.94] | 10.8 ± 6.1 [4.0-27.0] |
| **Case classification** |  |  |
| Imported (n=359) | 0.75 ± 0.15 [0.51-0.94] | 14.5 ± 7.0 [5.0-30.0] |
| Local<sup>a</sup> (n=215) | 0.75 ± 0.14 [0.49-0.91] | 13.2 ± 6.2 [4.0-25.0] |

Note: $A$ = number of unique alleles per locus; $H_E$ = expected heterozygosity; RACD = reactive case detection
<sup>a</sup> Local infections include intraport infections (n=12) and indigenous cases (n=203).

### Genetic diversity of parasites in Eswatini is similar to high transmission countries

To assess the relative genetic diversity of Eswatini to that of other malaria-endemic countries, estimates derived from Eswatini cases were compared to other genotyping studies using an overlapping subset of microsatellite markers. Our results demonstrate that mean H_E_ in Eswatini was significantly and consistently higher than those calculated among regions with similarly low malaria transmission intensities (e.g. Thailand [20], Vietnam [37], Honduras [38], and Indonesia [39]) and was more comparable to that of high transmission countries (e.g. Guinea [16], The Gambia [14], and Mali [15]) (Table 3). Comparison of mean MOI across countries showed that estimates from Eswatini were consistently comparable to estimates obtained from high transmission areas including in Ghana [40], Mali [17], and Burkina Faso [17] (Table 4). Among the countries listed in Table 4, the proportion of polyclonal infections was positively correlated with malaria incidence, with the exception of Eswatini, where the proportion of polyclonal infections was higher than all other comparator countries despite having very low malaria incidence.

**Table 3.** Comparison of mean expected heterozygosity (H_E_) observed in Eswatini isolates to previously published studies.

| Country | Year | Annual<br>confirmed<br>cases per 1000<sup>a</sup> | $H_E$ | | # of loci<br>compared | p-value<sup>c</sup> | References |
| --- | --- | --- | --- | --- | --- | --- | --- |
|  |  |  | Country | Eswatini<sup>b</sup> |  |  |  |
| Nicaragua | 2011 | 0.1 | 0.47 | 0.88 | 3 | 0.003 | Larranga et al (2013) [38] |
| Thailand | 2002-07 | 0.5 | 0.64 | 0.86 | 6 | 0.023 | Pumpaibool et al (2009) [20] |
| Vietnam | 1996-97 | 0.75 | 0.79 | 0.88 | 6 | 0.026 | Ferreira et al (2002) [37] |
| Honduras | 2011 | 1 | 0.31 | 0.88 | 3 | 0.017 | Larranga et al (2013) [38] |
| Indonesia (Papua) | 2011-14 | 1.5 | 0.69 | 0.87 | 8 | 0.02 | Pava et al (2017) [39] |
| Peru | 2003-07 | 3 | 0.53 | 0.88 | 3 | 0.089 | Branch et al (2011) [41] |
| Republic of Guinea | 2011 | 8 | 0.86 | 0.87 | 7 | 0.22 | Murray et al (2016) [16] |
| The Gambia | 2005-09 | 39 | 0.83 | 0.87 | 7 | 0.076 | Mobegi et al (2012) [14] |
| Mali | 2012-15 | 70 | 0.84 | 0.87 | 5 | 0.36 | Nabet et al (2016) [15] |

Note: $H_E$ = expected heterozygosity
<sup>a</sup> Estimates from the World Health Organization Malaria Report. 2000 incidence estimates are reported for studies conducted before 2000. Malaria incidence in Eswatini between 2014-2016 was estimated at 1 case per 1000 person-years.
<sup>b</sup> $H_E$ recalculated using only identical loci used in both studies
<sup>c</sup> p-value generated using a two-tailed Student's t-test comparing mean $H_E$ values across the subset of loci compared between studies.

**Table 4.** Comparison of the proportion of polyclonal infections and multiplicity of infection (MOI) of Eswatini isolates to previously published studies.

| Country | Year | Annual confirmed cases<br>per 1000<sup>a</sup> | Mean MOI | Polyclonal<br>infections (%) | References |
| --- | --- | --- | --- | --- | --- |
| Thailand | 2002-07 | 0.5 | -- | 0.28 | Ferreira et al (2002) [37] |
| **Eswatini** | **2014-16** | **1** | **2.2<sup>b</sup>** | **0.67** | **(this study)** |
| Indonesia (Papua) | 2011-14 | 1.5 | 1.2<sup>c</sup> | 0.17 | Pava et al (2017) [39] |
| Peru | 2003-07 | 3 | -- | 0.34 | Branch et al (2011) [41] |
| Ghana | 2003-04 | 28 | 2.3<sup>d</sup> | -- | Agyeman-Budu (2013) [40] |
| Mali | 2012-15 | 70 | 2.6<sup>c</sup> | 0.59 | Auburn et al (2012) [17] |
| Burkina Faso | 2012 | 248 | 3.0<sup>c</sup> | -- | Auburn et al (2012) [17] |
| Southern Mozambique<br>(Maputo Province) | 1997-99 | 312 | 1.8<sup>d</sup> | 0.49 | Mayor et al (2003) [42] |

Note: MOI = multiplicity of infection
<sup>a</sup> Estimates from the World Health Organization Malaria Report. 2000 incidence estimates are reported for studies conducted before 2000. Malaria incidence in Eswatini between 2014-2016 was estimated at 1 cases per 1000 person-years.
<sup>b</sup> Calculated from the second highest MOI.
<sup>c</sup> Calculated from maximum alleles found at any one microsatellite.
<sup>d</sup> Calculated from *msp-2* typing.

### Imported cases are more complex than local cases

Global measures of population-level and within-host genetic diversity were compared between imported and locally-acquired cases, as determined by travel history. Population-level diversity was similar between imported and locally-acquired cases (Table 2), with little evidence of genetic differentiation (G’_ST_=0.013). Despite the similarities observed in population-level genetic diversity, infections among locally-acquired cases were less complex than imported cases (i.e. exhibited lower within-host diversity). A larger proportion of local cases were monoclonal infections compared to imported cases (41% versus 29%; p=0.004), and had a slightly lower mean MOI (2.0 vs. 2.4; p=0.004) (Table 1, Figure 3). A similar relationship was observed in measures of F_WS_. Mean F_WS_ was significantly higher among local cases compared to imported cases (0.91 vs. 0.85; p=0.03), and a higher proportion of local cases were considered clonal infections by this measure, that is, 44% of local cases had a mean F_WS_ greater than 0.95 compared to 34% of imported cases (p=0.018). Sensitivity analyses using reweighted estimates to account for potential bias in which samples were genotyped showed similar patterns of within-host diversity between imported and local cases (Supplementary Table 4).

**Figure 3.**
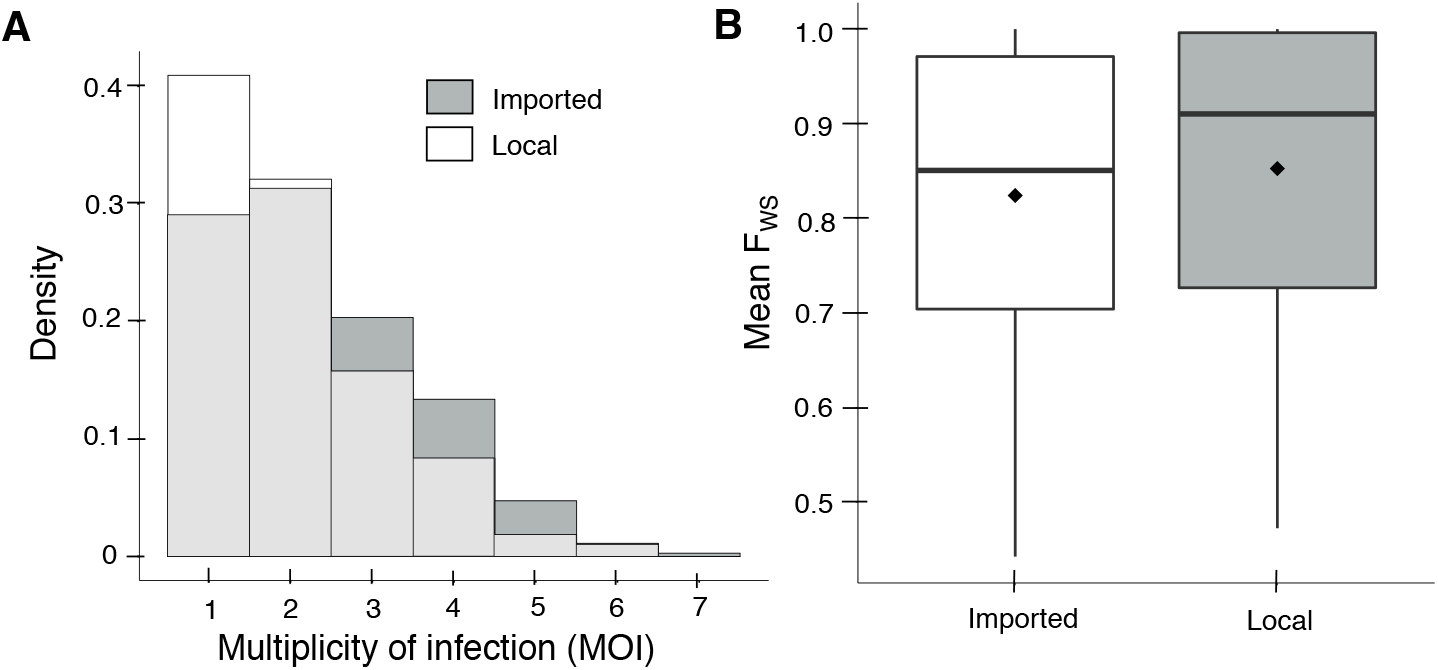
Distribution of multiplicity of infection (MOI) (A) and mean F_WS_ (B) by imported and locally-acquired cases. In both figures, white indicates the distribution of MOI and F_WS_ among imported cases and dark grey indicates distribution among locally-acquired cases. In Figure 3A, light grey represents the overlap between imported and locally-acquired cases. In Figure 3B, the horizontal line indicates the median, the diamond marks the mean, and the upper and lower bounds of the box indicate the 25^th^ and 75^th^ quartiles.

### Inclusion of MOI and F_WS_ does not meaningfully improve discrimination between imported and local cases

Given that there were significant differences in within-host diversity metrics between local and imported cases, we evaluated whether these metrics could be used to determine whether a case was local versus imported, since travel history data are not always available or accurate. Compared to a base model that included district of residence, season, whether a case was passively or reactively detected, age, gender, and occupation to predict malaria importation, the model that included F_WS_ and MOI marginally improved the goodness of fit (difference in the log-likelihood chi-squared statistic=+8.17; p=0.017), but had negligible effects on improving the discrimination between imported and local cases (difference in AUC=+0.002; p=0.6) (Supplementary Figure 3).

## DISCUSSION

Despite sustaining low malaria transmission for nearly a decade, our study demonstrated that malaria in Eswatini is largely characterized by polyclonal, complex *P. falciparum* infections and a markedly diverse parasite population. This evidence was generated by genotyping two-thirds of all documented, symptomatic and asymptomatic *P. falciparum* infections in the country over the span of two years, thus represents a comprehensive picture of the parasite population. The genetic diversity and large effective population size reveal a *P. falciparum* population similar to those observed among high transmission regions of sub-Saharan Africa [14-17, 42]. Importantly, imported and local cases had similar levels of population-level diversity, and though MOI and F_WS_ were higher among imported cases, the magnitude of these differences did not allow for accurate discrimination between these two groups.

These data present an important caveat to the “genomic thermometer” framework: namely, the assumption that as malaria reaches very low levels of transmission, *P. falciparum* populations are expected to consistently exhibit low within-host diversity, low heterozygosity, a small effective population size, and genetically fragmented subpopulations [8, 11, 18-22]. Although a monotonic relationship between diversity and transmission may be expected in isolated settings where local transmission is low, resulting in genetic bottlenecking of local parasites [11, 13, 18, 19, 22, 43], and importation is uncommon or is imported from areas with low genetic diversity [21], this framework fails to accurately represent countries like Eswatini, where low malaria transmission has been sustained yet the parasite population remains highly diverse. Though this study contradicts previous studies from low transmission settings outside of sub-Saharan Africa, this apparent paradox between genetic diversity and transmission intensity is not surprising. Indeed, this finding is consistent with the known epidemiological patterns of malaria in Eswatini, where *P. falciparum* transmission needs to be considered in the context of substantial human mobility between Eswatini and neighboring high transmission countries [3, 4, 7], which has resulted in the frequent importation of highly diverse parasites into a setting of limited local transmission [44, 45].

In this setting, further decreases in local transmission may have actually resulted in increases in parasite diversity due to: (1) an increase in the proportion of imported infections, which tend to have higher parasite diversity, and (2) shorter chains of local transmission. Shorter chains would be expected to increase population diversity, since there are fewer related parasites circulating, and to increase within-host diversity, since infections, on average, maintain higher within-host diversity as decreases from those of imported infections would be expected to occur with each generation of local transmission in the absence of any appreciable superinfection. As more sub-Saharan African countries with similar challenges move towards malaria elimination, this apparent paradox may become the rule rather than the exception, and as such, these metrics of diversity may not be the most appropriate methods to assess changes in transmission intensity in these settings.

Malaria importation remains a significant barrier for eliminating malaria in many countries and is further complicated by the difficulty in accurately distinguishing between imported and locally-acquired infections. In Eswatini, imported and local infections had similar levels of population genetic diversity, with little evidence of genetic differentiation between groups. Imported cases did display significantly higher within-host diversity compared to local cases, but these differences were minimal, likely due to the limited length of local transmission chains, as described above. As a result, these differences were too small to be used to accurately discriminate between imported and local infections.

While one of the most comprehensive studies of its kind to date, analyses of genotyped cases may not have been a representative sample of all infections present in the country. Though the study was able to successfully genotype 67% of all documented cases and included sensitivity analyses confirming within-host diversity measures produced generalizable results, it is possible that undocumented (e.g. asymptomatic) infections were substantially different from those detected. Fortunately, this study was able to evaluate 80 cases captured through reactive case detection, which demonstrated no differences with symptomatic cases. Second, though assignment of imported and local infections were conducted by surveillance officers using detailed travel history information and demographic factors, and later rechecked by study team members, its possible that cases may have been misclassified. This may have biased the discriminatory capacity of MOI towards the null.

In conclusion, these findings demonstrate that *P. falciparum* genetic diversity and transmission intensity do not always follow a simple, monotonic relationship and suggests that the *P. falciparum* parasite population in an eliminating country with strong connectivity to high transmission countries is likely to be highly diverse. Thus, a more nuanced approach must be taken when interpreting the results from genetic analyses, particularly as a means of monitoring and evaluating the impact of malaria interventions in these settings. In countries such as Eswatini, where transmission is largely dominated by the importation of cases from higher transmission countries, traditional metrics derived from genotyping data, such as MOI, may not be informative. In these settings, detailed characterization of *P. falciparum* transmission, or interruption thereof, may be better achieved through a more comprehensive analysis of transmission networks, including accounting for infections not observed through the national malaria surveillance database.

## ACKNOWLEDGEMENTS

The authors would like to thank the participants for contributing their samples to the study; the health worker staff who performed sample and data collection; and the Eswatini National Malaria Program and the Clinton Health Access Initiative for their support in data collection.

