## Supplementary Appendix for "High genetic diversity of *Plasmodium falciparum* in the low transmission setting of the Kingdom of Eswatini"

Supplementary Tables and Figures

**Supplementary Table 1.** Characteristics of genotyped and non-genotyped *Plasmodium falciparum* cases detected in Eswatini between July 2014 and June 2016.

| Characteristics | All cases (n=880) | Genotyped (n=582) | Not genotyped* (n=298) | p-value** |
|---|---|---|---|---|
| ***Detection method*** | | | | <0.0001 |
| Passive surveillance | 725 (82.4) | 502 (86.3) | 223 (74.8) | |
| RACD | 155 (17.6) | 80 (13.7) | 75 (25.2) | |
| ***Season*** | | | | <0.0001 |
| 2014-2015 | 635 (72.2) | 389 (66.8) | 246 (82.6) | |
| 2015-2016 | 245 (27.8) | 193 (33.2) | 52 (17.4) | |
| ***District*** | | | | <0.0001 |
| Hhohho | 157 (19.8) | 115 (22.4) | 42 (14.5) | |
| Lubombo | 389 (48.5) | 235 (45.8) | 154 (53.3) | |
| Manzini | 239 (29.8) | 158 (30.8) | 81 (28.0) | |
| Shiselweni | 17 (2.1) | 5 (1.0) | 12 (4.2) | |
| ***Case classification*** | | | | <0.0001 |
| Imported | 485 (55.1) | 359 (61.7) | 126 (42.3) | |
| Locally acquired | 376 (42.7) | 215 (36.9) | 161 (54.0) | |
| Unknown | 19 (2.2) | 8 (1.4) | 11 (3.7) | |
| ***Gender*** | | | | 0.0088 |
| Female | 222 (27.2) | 128 (24.2) | 94 (33.0) | |
| Male | 593 (72.8) | 402 (75.8) | 191 (67.0) | |
| ***Age, median (IQR)*** | 25.3 (11.3-37.1) | 26.6 (11.9-37.1) | 23.9 (9.8-37.2) | 0.31 |
| ***Occupation*** | | | | 0.0003 |
| Child | 135 (15.6) | 75 (13.1) | 60 (20.5) | |
| Farming/Agriculture | 93 (10.8) | 64 (11.2) | 29 (9.9) | |
| Manual Laborer | 103 (11.8) | 77 (13.5) | 25 (8.6) | |
| Manufacturing/Factory | 61 (7.1) | 52 (9.1) | 9 (3.1) | |
| Small-market trader | 66 (7.6) | 50 (8.7) | 16 (5.5) | |
| Student | 186 (21.5) | 127 (22.0) | 59 (20.2) | |
| Unemployed | 170 (19.7) | 97 (17.0) | 73 (25.0) | |
| Other | 48 (5.9) | 30 (5.2) | 21 (7.2) | |

Col = column; RACD = reactive case detection; IQR = interquartile range

Note: Values may not add up to total N due to missing values. Values for categorical variables represent N (Column %) and continuous variables show median (IQR).

[a] Non-genotyped samples include genotype failures, samples where dried blood samples were not collected, and may also include samples that were falsely positive by rapid diagnostic test or microscopy and thus were not truly cases.

[b] p-value generated using two-tailed Chi-squared test for categorical variables or Student's t-test for continuous variables.

**Supplementary Figure 1.** Density plot of propensity scores (e.g. estimated probabilities of being genotyped) stratified by actually observed genotyped and non-genotyped cases.

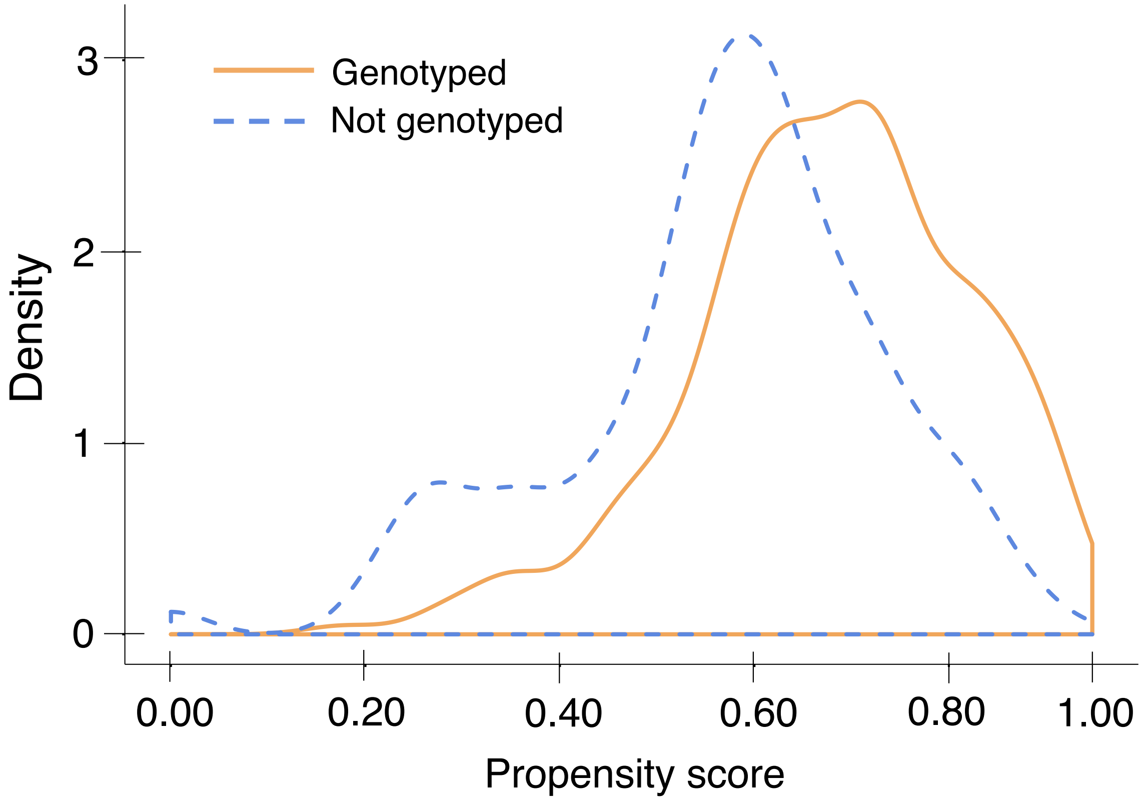

**Supplementary Figure 2.** Distribution of mean $F_{WS}$ by MOI among genotyped Eswatini samples.

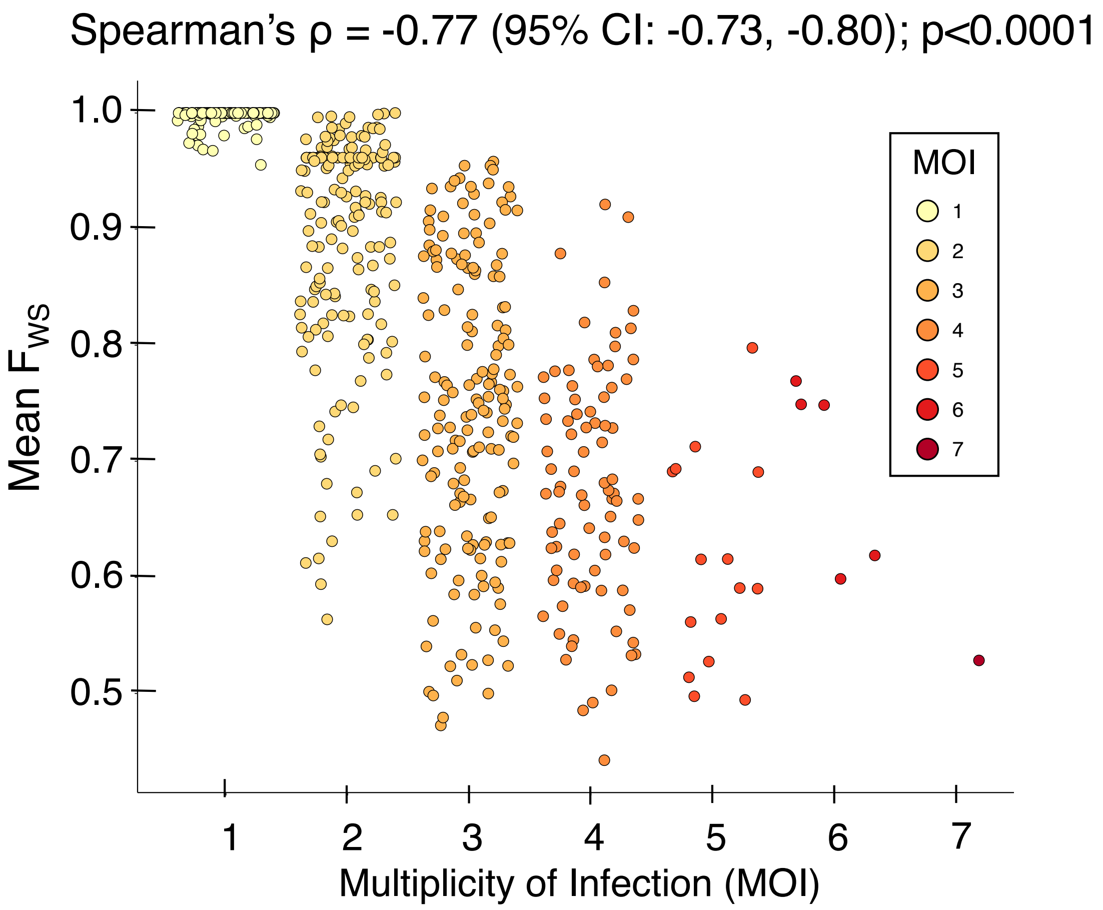

**Supplementary Table 2.** Expected Heterozygosity ($H_E$) Assessed at 26 Microsatellites.

| Locus | Total | Case classification | |
|---|---|---|---|
| | | Imported | Local |
| Ara2 | 0.862 | 0.864 | 0.848 |
| AS1 | 0.546 | 0.533 | 0.558 |
| AS2 | 0.770 | 0.787 | 0.743 |
| AS3 | 0.858 | 0.861 | 0.858 |
| AS7 | 0.533 | 0.511 | 0.600 |
| AS8 | 0.572 | 0.547 | 0.605 |
| AS11 | 0.715 | 0.711 | 0.726 |
| AS12 | 0.528 | 0.511 | 0.544 |
| AS14 | 0.773 | 0.779 | 0.764 |
| AS15 | 0.848 | 0.839 | 0.854 |
| AS19 | 0.668 | 0.663 | 0.675 |
| AS21 | 0.520 | 0.529 | 0.492 |
| AS25 | 0.864 | 0.859 | 0.864 |
| AS31 | 0.860 | 0.860 | 0.860 |
| AS32 | 0.718 | 0.686 | 0.722 |
| AS34 | 0.551 | 0.563 | 0.526 |
| B7M19 | 0.604 | 0.605 | 0.605 |
| PFG377 | 0.655 | 0.678 | 0.606 |
| PfPK2 | 0.919 | 0.924 | 0.910 |
| PolyA | 0.931 | 0.935 | 0.905 |
| TA1 | 0.897 | 0.894 | 0.883 |
| TA40 | 0.898 | 0.896 | 0.910 |
| TA60 | 0.827 | 0.830 | 0.818 |
| TA81 | 0.857 | 0.853 | 0.864 |
| TA87 | 0.883 | 0.883 | 0.877 |
| TA109 | 0.818 | 0.820 | 0.824 |
| **Mean $H_E$ ± SD** | 0.75 ± 0.14 | 0.75 ± 0.15 | 0.75 ± 0.14 |

**Supplementary Table 3.** The Number of Unique Alleles (A) Assessed at 26 Microsatellite Loci.

| Locus | Total | Case Classification | |
|---|---|---|---|
| | | Imported | Local |
| Ara2 | 13 | 13 | 13 |
| AS1 | 7 | 7 | 6 |
| AS2 | 11 | 11 | 11 |
| AS3 | 11 | 11 | 11 |
| AS7 | 13 | 11 | 13 |
| AS8 | 7 | 5 | 6 |
| AS11 | 14 | 14 | 10 |
| AS12 | 8 | 6 | 6 |
| AS14 | 15 | 14 | 15 |
| AS15 | 14 | 14 | 13 |
| AS19 | 17 | 16 | 15 |
| AS21 | 9 | 7 | 5 |
| AS25 | 23 | 21 | 20 |
| AS31 | 24 | 24 | 19 |
| AS32 | 21 | 18 | 16 |
| AS34 | 8 | 7 | 4 |
| B7M19 | 7 | 7 | 5 |
| PFG377 | 7 | 7 | 7 |
| PfPK2 | 22 | 21 | 22 |
| PolyA | 32 | 30 | 25 |
| TA1 | 29 | 29 | 23 |
| TA40 | 24 | 22 | 23 |
| TA60 | 12 | 11 | 10 |
| TA81 | 16 | 16 | 14 |
| TA87 | 17 | 16 | 15 |
| TA109 | 20 | 18 | 15 |
| **Mean A ± SD** | 15.4 ± 7.1 | 14.5 ± 7.0 | 13.2 ± 6.2 |

**Supplementary Table 4.** Propensity-score weighted MOI and mean $F_{WS}$ from sensitivity analyses

| Measures of within-host diversity | Total | Case classification | | |
|---|---|---|---|---|
| | | Imported | Local | p-value[a] |
| % polyclonal infections (95% CI) | 75 (72-79) | 78 (71-80) | 70 (63-76) | 0.032 |
| Mean MOI ± SD | 2.4 ± 1.2 | 2.5 ± 1.2 | 2.3 ± 1.1 | 0.19 |
| Mean $F_{WS}$ ± SD | 0.838 ± 0.154 | 0.83 ± 0.24 | 0.86 ± 0.20 | 0.036 |

[a] p-values calculated from simple linear regression with probability weights.

**Supplementary Figure 3.** Receiver operating characteristic (ROC) curves of case classification models. Blue line indicates ROC curve of model with MOI and $F_{WS}$ as covariates. Orange line indicates ROC curve of model with only epidemiological (epi) covariates. Green line indicates ROC curve of full model with both epi and genetic covariates. The null hypothesis (black dashed line) is that the area under the curve (AUC) equals 0.5.

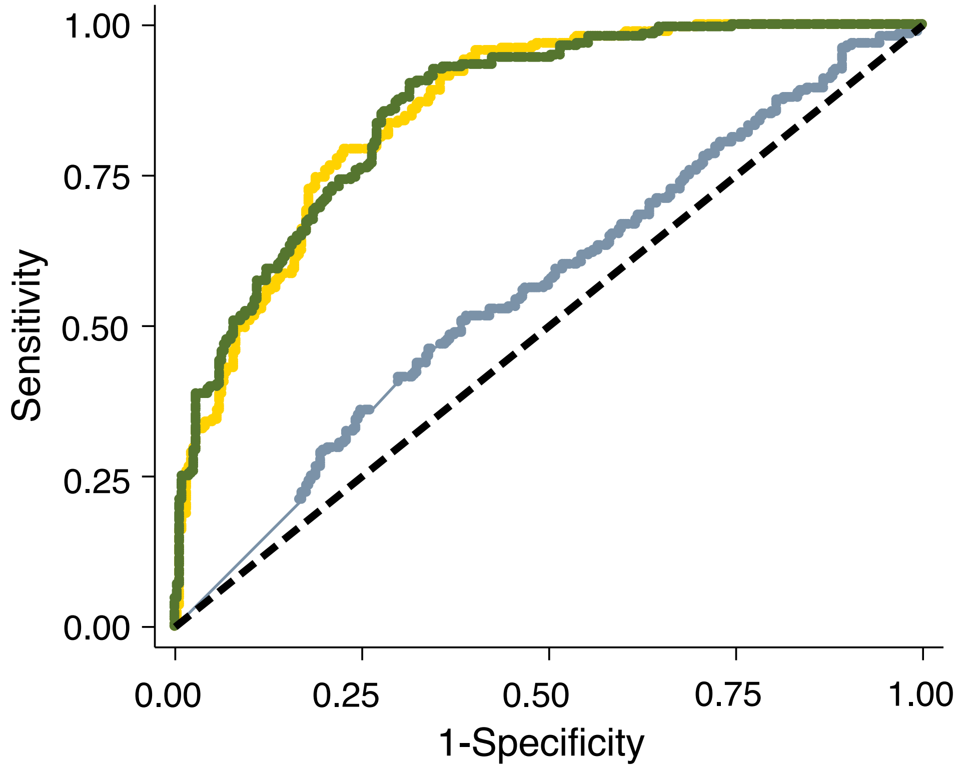
